# Disrupting inferior frontal cortex activity alters affect decoding efficiency from clear but not from ambiguous affective speech

**DOI:** 10.1101/2021.12.15.472758

**Authors:** Leonardo Ceravolo, Marius Moisa, Didier Grandjean, Christian Ruff, Sascha Frühholz

## Abstract

The evaluation of socio-affective sound information is accomplished by the primate auditory cortex in collaboration with limbic and inferior frontal cortex (IFC)—often observed during affective voice classification. Partly opposing views have been proposed, with IFC either coding cognitive processing challenges in case of sensory ambiguity or representing categorical object and affect information for clear voices. Here, we presented clear and ambiguous affective speech to two groups of human participants during neuroimaging, while in one group we inhibited right IFC activity with transcranial magnetic stimulation. IFC activity inhibition led to faster affective decisions, more accurate choice probabilities, reduced auditory cortical activity and increased fronto-limbic connectivity for clear affective speech. This indicates a more intermediate functional property of the IFC than assumed—namely with normal activity representing a more deliberate form of affective sound processing (i.e., enforcing cognitive analysis) that flags categorical sound decisions with precaution (i.e., representation of categorical uncertainty).

**Teaser:** Inferior frontal cortex enforces cognitive analyses during affect decisions with different levels of sensory ambiguity.

## Introduction

The neural processing and classification of acoustic information involves a cortical neural network beyond auditory decoding in the auditory cortex (AC; Staib and Frühholz 2020). The core of this neural network especially for decoding acoustic and social information conveyed by acoustic voice signals involves an integrated functioning of the AC together with various regions in the inferior frontal cortex (IFC; Rauschecker and Scott 2009, Roswandowitz, Swanborough et al. 2020). This auditory-frontal network is especially relevant for extracting socio-affective information from voice signals (Frühholz and Grandjean 2012, Frühholz and Grandjean 2013, Dricu, Ceravolo et al. 2017, Swanborough, Staib et al. 2020), such as decoding emotional information expressed in affective voices and affective intonations in speech. In the latter case, the auditory-frontal network is often accompanied by neural activity in the limbic system, and this limbic activity is primarily located in the amygdala (Frühholz, Trost et al. 2014, Frühholz, Trost et al. 2016).

Within this broad auditory-frontal-limbic network for voice signals processing and voice information decoding (Dricu, Ceravolo et al. 2017, Dricu and Frühholz 2020, Frühholz and Schweinberger 2020), the involvement of the IFC in decoding emotions from affective speech has been consistently reported (Frühholz and Schweinberger 2020, Roswandowitz, Swanborough et al. 2020, Swanborough, Staib et al. 2020), but its functional role in the neural network and its functional contribution to voice signal processing and especially in vocal affect decoding remained debated. Some previous studies indicated that social affective decoding from voice seemed restricted to the AC (Grandjean, Sander et al. 2005, Ethofer, Van De Ville et al. 2009) without any functional relevance of the IFC. However, more recent studies highlighted the functional relevance of the IFC in two possible directions. First, some studies seem to suggest that the IFC is especially relevant for elaborate evaluations and classification of socio-affective information in voices (Schirmer, Zysset et al. 2004, Fecteau, Armony et al. 2005). These processes are supposed to happen down-stream to the acoustical analysis in the AC (Rauschecker and Scott 2009, Frühholz, Ceravolo et al. 2012, Frühholz and Grandjean 2012) with different IFC subregions coding for the complexity of the evaluation and classification task (Frühholz and Grandjean 2013, Dricu, Ceravolo et al. 2017, Dricu and Frühholz 2020). According to these studies, the IFC would represent the rather domain-general task, evaluation, and classification difficulty, and would show increased activity for any challenging affective evaluation and classification tasks. This notion would support the related view on the IFC as a region that exerts some top-down control on the AC depending on certain task requirements (Dricu, Ceravolo et al. 2017, Roswandowitz, Swanborough et al. 2020) and explicit sound classifications demands (Roswandowitz, Swanborough et al. 2020). The latter seems especially relevant when classifying ambiguous voices and affect information (Bestelmeyer, Maurage et al. 2014), which requires that acoustical analyses in AC are supplemented by functional support by neural processing resources of the IFC (Swanborough, Staib et al. 2020). In fact, ambiguous voices roughly contain the same acoustic information as non-ambiguous voices yet they are perceived differently (Hoekert, Bais et al. 2008, Hoekert, Vingerhoets et al. 2010).

Unlike the aforementioned evidence, some other studies especially from animal research have shown a more concrete psychoacoustic role of the IFC instead of representing abstract decisional task demands, as some IFC subregions seem to be directly involved in the perceptual discrimination of different types of conspecific vocalizations (Averbeck and Romanski 2006, Cohen, Russ et al. 2009, Frühholz and Grandjean 2013). According to these studies, the IFC would function pre-dominantly as a domain-specific higher-order auditory processing node that represents sound information in relation to categorical certainty or discriminability (Frühholz and Schweinberger 2020), especially during explicit voice signal classification tasks (Roswandowitz, Swanborough et al. 2020). This notion would suggest some kind of opposite IFC activity patterns as suggested by the first line of studies. Especially, this notion would predict higher IFC activity for clear as opposed to ambiguous affective voices, as clear voices would allow a more direct categorical representation. Given these two different hypothesis about the role of the IFC in processing affective voices, more knowledge is critical to determine the functional role of the IFC for the fine-grained classification of affective voices at different levels of acoustic ambiguity.

To mechanistically investigate the functional role of the IFC in the classification of socio-affective voice information, an experimentally induced alteration of IFC activity especially during the processing of both clear and ambiguous affective speech would reveal direct evidence. Unlike clear affective expressions in voices which represent categorical certainty as predicted by the second hypothesis, ambiguous vocal affect might trigger computations in the IFC given their challenging perceptual and cognitive processing according to the first hypothesis. Both cases might additionally require integrated IFC-STC functioning in terms of neural connectivity (Roswandowitz, Swanborough et al. 2020), either for top-down IFC-to-STC facilitations or as co-representations of categorical affective information (Steiner, Bobin et al. 2021). The use of transcranial magnetic stimulation (TMS) appears to be a reliable and non-invasive option to especially inhibit a proper functioning of computations performed in the IFC in this context. Continuous theta burst stimulation (cTBS), a patterned pulse stimulation, as well as repetitive TMS (rTMS) are known to create reversible neural activity alteration (Oberman, Edwards et al. 2011). Although the neural effects induced by the cTBS procedure are much faster and potentially longer-lasting than those of rTMS, the cTBS procedure was rather scarcely used for social and especially voice processing studies, even less in combination with functional magnetic resonance imaging (fMRI). Previous studies using cTBS procedure revealed impairment on voice identity but not on affect discrimination tasks when applied over the premotor cortex (Banissy, Sauter et al. 2010). Similar results were observed in a combined cTBS-fMRI study targeting the premotor cortex in vocal affect processing, showing that cTBS triggered differential brain patterns in fronto-parietal, parahippocampal, and IFC brain regions (Agnew, Banissy et al. 2018).

Compared to cTBS, the use of rTMS revealed more mixed results regarding socio-affective processing from voices. First, the rTMS procedure was unsuccessful on both the superior temporal cortex (STC; Jiahui, Garrido et al. 2017) and IFC (Hoekert, Vingerhoets et al. 2010) as it did not lead to expected behavioral changes, for example, in terms of a difficulty to classify vocal affect. Second, some reaction time differences were observed for the affect classifications of clear sad and happy voices by the use of rTMS over the right but not the left STC (Alba-Ferrara, Ellison et al. 2012), showing longer delays to respond in the classification task. Similar results were observed as well when rTMS was applied over the right premotor cortex (Sammler, Grosbras et al. 2015) and over the right anterior STC and fronto-parietal operculum (Hoekert, Bais et al. 2008). Third, again no behavioral differences were observed in yet another study when rTMS was applied to either the bilateral STC or IFC (Jacob, Brück et al. 2014), even though abovementioned results tell another story, and even though the right STC was shown to causally interact with voice detection, independently of emotion (Bestelmeyer, Belin et al. 2011). Some methodological variations may explain these inconsistencies, most importantly regarding the type of TMS procedure (rTMS or cTBS) and especially large variations in IFC target region(s).

Given these limitations and inconsistencies concerning socio-affective classification from affective speech with inhibitory effects especially applied to the IFC, we designed a three-alternative classification task on clear and ambiguous vocal affect while brain activity was influenced and recorded in a combined cTBS-fMRI setup. In order to address more precisely the cognitive and evaluative role of the IFC in such contexts (Schirmer and Kotz 2006), we created stimuli containing a gradual blending of angry and fearful voices to manipulate affective ambiguity expressed in these voices. Anger and fear are distinctive and well-recognized emotions expressed in vocal affect, and a blending of anger and fear leads to considerable affective ambiguity given their opposite nature for behavioral adaptations in listeners (Whiting, Kotz et al. 2020). This setup also allowed to determine brain activity patterns as well as functional connectivity changes following the cTBS procedure between two groups of participants in a between-group design. In one group, cTBS was applied to the right IFC (experimental group, ‘EG’), while in another group cTBS was applied to the vertex as a control brain site (control group, ‘CG’). We hypothesized that: (a) cTBS over the IFC would alter affect classifications and response speed especially for ambiguous affective voices—according to challenging perceptual and cognitive processing of these voices (Schirmer and Kotz 2006, Hoekert, Vingerhoets et al. 2010, Frühholz and Grandjean 2012, Frühholz and Grandjean 2013)—but also potentially for clear affective voices—representing categorical certainty; (b) cTBS over the IFC should also lead to a distinct reorganization of brain activity patterns while classifying the most ambiguous affective voices, especially within the right AC/STC for voice processing (Frühholz, Ceravolo et al. 2012, Frühholz and Grandjean 2012, Frühholz and Schweinberger 2020) and in the limbic system (i.e., amygdala) for affect processing (Frühholz, Trost et al. 2014); and (c) we expected altered functional coupling between IFC and the limbic and auditory cortical system during cTBS to the IFC given the important status of the IFC in the neural network for socio-affective voice processing (Frühholz, Ceravolo et al. 2012, Frühholz and Grandjean 2012, Grandjean 2020).

## Results

The present study used a combined cTBS-fMRI setup to non-invasively and temporarily alter brain functioning in the right IFC while human participants of the EG (cTBS over the right IFC; Fig.2) and the CG (cTBS over the vertex; Fig.2) performed a three-alternative forced choice task on clear and ambiguous affective speech. We note here that all *p*-values reported for the statistical tests on the behavioral choice and reaction time data are corrected for multiple comparisons using the Bonferroni method.

For the affective forced choice task, we created blends of vocally expressed anger and fear using a voice morphing procedure (Rauschecker and Scott 2009, Staib and Frühholz 2020) on a set of commonly available and validated vocal affective bursts, namely the Montreal affective voice database or ‘MAV’ (Belin, Fillion-Bilodeau et al. 2008). From this database we selected five male and five female voices. We created blends of affective voices ranging from 100% of one emotion to 100% of the other emotions in steps of 10% morphing, resulting in 11 different stimulus categories (Fig.1**C**): 60-100% fear proportions (F_60_, F_70_, F_80_, F_90_, F_100_), 60-100% anger proportions (A_60_, A_70_, A_80_, A_90_, A_100_), and the 50/50 mix of anger and fear in the condition AF_50_.

**Fig.1:**
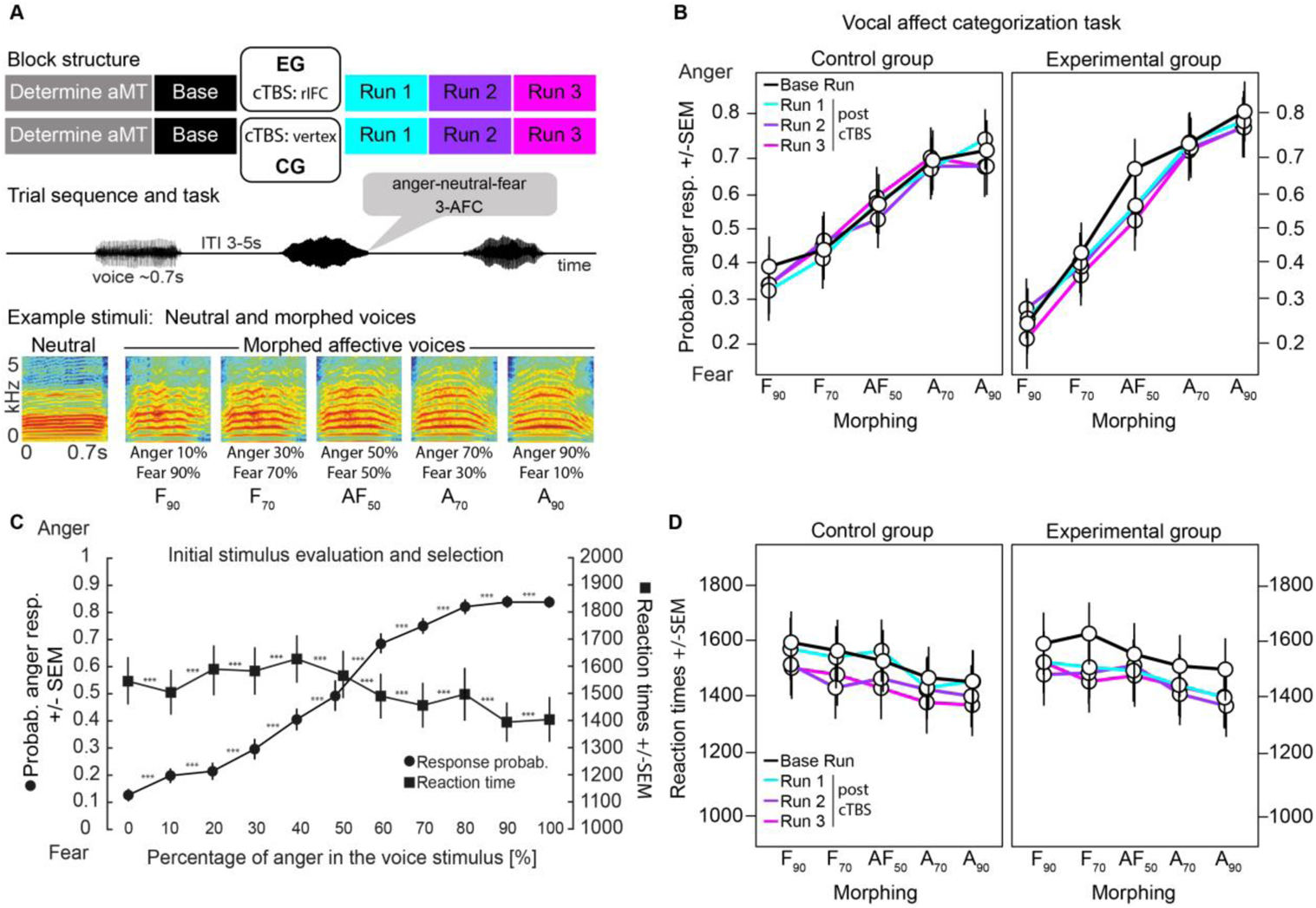
Experimental timeline, example affective voices, and behavioral results for initial stimulus selection and the vocal affect categorization task. (**A**) Experimental timeline for one participant, showing the different procedures for each group and detailing the three-alternative forced choice task (3-AFC) with examples of stimuli including spectrograms of neutral and morphed voices with percentage of each morphed emotion (anger, fear). (**B**) Probability of an ‘anger’ response as a function of the morphing procedure for each run, per group for the vocal affect categorization task. (**C**) Following the initial morphing procedure, we had an independent sample of nineteen right-handed participants (10 female, 9 male, mean age 32y, SD 5.2) evaluate the emotionally blended stimuli. Participants were asked to categorize each stimulus (n=110; 10 trials for each of the 11 morphing levels) by a keypress (1: the voice expresses fear; 2: the voice epxresses anger; the keys were counterbalanced across participants). The line plots illustrate on the *y* axis the probability of an ‘anger’ response (emotion categorization response; black circles) and reaction times (black squares) according to each morphing level (*x* axis) with errorbars representing the standard error of the mean (SEM). (**D**) Averaged reaction times results for the vocal affect categorization task, for each morphing level, run and group. Error bars represent the standard error of the mean (SEM). *aMT* active motor threshold; *cTBS* continuous theta-burst stimulation (transcranial magnetic stimulation procedure); *IFC* inferior frontal cortex target region (MNI *xyz* 54 34 10) based on *a priori* data (see TMS site localization section); *CG* control group; *EG* experimental group; *ITI* inter-trial interval; *cTBS* continuous theta burst stimulation; *Probab* probability; *resp* response. \*\*\**p*<.0001.

For selecting the appropriate stimuli and morphing rates for the main experiment, we asked an independent sample of participants (N=19) to classify these stimuli in a 2AFC task as portraying either ‘anger’ or ‘fear’. Our aim was to select one emotionally ambiguous morphing level as well as four other gradually less ambiguous emotional voices for a total of five morphed voices. Using generalized linear mixed-effects (‘glmer’) modelling with the participants’ choices as dependent variable, illustrating the probability of an ‘anger’ response. We observed a significant main effect of morphing on the categorization (χ^2^(10)=1094.3, *p*<.0001); reaction time data were of no major interest here but are reported in Fig.1**C** and Table S1. We used planned comparisons to explore which morphing levels would be more suitable—i.e., accurately evaluated—and contrasts showed significant differences between any pair of morphing levels (all *p*<.0001, see Fig.1, Table S2). Since the AF_50_ condition embodied the highest affective ambuguity, we selected it as well as two less ambiguous morphing levels per emotion in equal steps of 20% morphing, that is A_90_, A_70_, F_90_ and F_70_. This model explained 40.25% of the variance including both fixed and random effects (R2c=0. 4025) and 30.46% of the variance solely for fixed effects (R2m=0.3046).

According to this selection criteria, the stimuli for the main experiment consisted of expressions of vocal affect containing 10%, 30%, 50%, 70%, or 90% of fear and therefore 90%, 70%, 50%, 30%, and 10% of anger at the same time, respectively (labels: A_90_, A_70_, AF_50_, F_70_, F_90_; Fig.1**A**). The 10% conditions referred to rather clearly expressed vocal affect, the 50% condition was of highest affective ambiguity, and the 30% morphing level referred to an intermediate level of ambiguity. In the main fMRI task, participants therefore listened to these five expressions of morphed vocal affect as well as to neutral vocalizations in the main experiment, and they were asked to classify them as ‘anger’, ‘fear’, or ‘neutral’. The neutral voices were added from each speaker and served as a control condition during affective decisions. Participants performed these decisions in one run before cTBS (cTBS_pre_), and in three runs after cTBS was applied (cTBS_post_) to the vertex (CG) or to the right IFC (EG) (see Fig.1**A**). The inclusion of a cTBS_pre_ run was motivated by the need for a control measure of brain activity between groups independently of the cTBS procedure, namely a sanity check for the main task providing an overview of brain activity and behavior depending merely on task conditions.

For the main fMRI task and for both behavioral and neuroimaging data analysis, the triple interaction between *group*(CG,EG), *morphing*(A_90_, A_70_, AF_50_, F_70_, F_90_) and *run*(cTBS_pre_ run,cTBS_post_ run1-3) was computed using two contrasts of interest: [CG > EG] x [A_90_, A_70_, F_90_, F_70_ > AF_50_] x [cTBS_post_ > cTBS_pre_] and its inverse [CG > EG] x [AF_50_ > A_90_, A_70_, F_90_, F_70_] x [cTBS_post_ > cTBS_pre_]. For the *morphing* factor in these contrasts, a specific contrast weighting was applied—namely [2*(A_90_) + 1*(A_70_) −6*(AF_50_) + 1*(F_70_) + 2*(F_90_)]—to better model and characterize the morphing percentages of the affective voices in the contrast vectors. Such weighting was then of course adapted to each comparison between *run* and *group* with necessary sign inversion(s) and multiplication(s).

### Right IFC as target region for cTBS procedure

As mentioned in the introduction, decisions on vocal affect are typically associated with activity in bilateral IFC. In the current study, we acquired the data of the CG first and we chose the right IFC, namely the right inferior frontal gyrus *pars triangularis* [IFGtri, MNI *xyz* 54,34,10; Fig.2**AB**], as target region for cTBS. This was based on the observation that the right IFGtri showed higher activity for clear as opposed to ambiguous voice processing in the CG especially when contrasting cTBS_post_ compared to cTBS_pre_ runs. This contrast matched also the pattern of contrasts we were subsequently interested in for to the data obtained for the EG.

**Fig.2:**
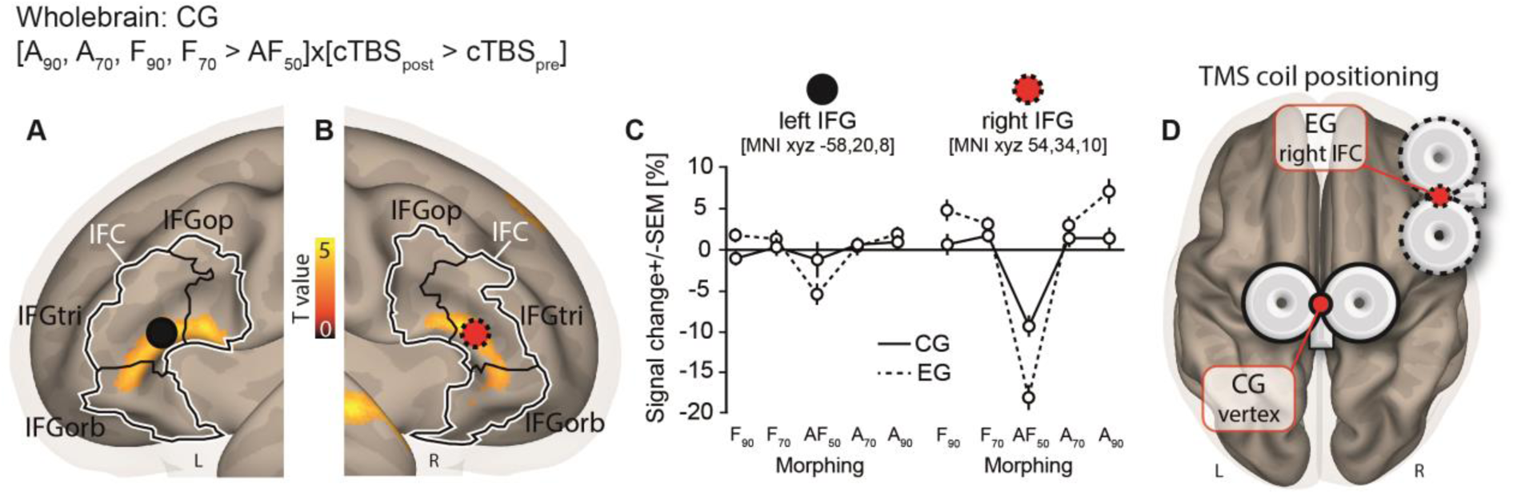
Neuroimaging results of the affect classification task for the CG with a focus on the IFC. (**A**) Whole-brain data showing control group (CG) activations (black circle: left IFC maxima) for the [A_90_, A_70_, F_90_, F_70_ > AF_50_] x [cTBS_post_ > cTBS_pre_] contrast with continuous theta burst stimulation (cTBS) over the vertex, left hemisphere. (**B**) Activations for the same contrast in the right hemisphere, showing global maxima activity in the right inferior frontal gyrus *pars triangularis* (red circle, dashed black outline), used as the target region for the cTBS procedure of the EG. (**C**) Percentage of signal change using a cube of 27 voxels adjacent to the peak voxel—extracted according to activity maxima of the [A_90_, A_70_, F_90_, F_70_ > AF_50_] x [cTBS_post_ > cTBS_pre_] contrast for the CG—for both groups in the left (MNI *xyz* −58, 20, 8; black circle) and right (MNI *xyz* 54,34,10; red circle, dashed black outline) IFGtri for each morphing condition. (**D**) TMS coil positioning for the cTBS procedure in both groups (CG: vertex, red circle with continuous black outline, MNI *xyz* [0, −20, 80]; EG: right IFC within the IFGtri, red circle with dashed black outline, MNI *xyz* [54, 34, 10]), decided after the CG was scanned in whole. Wholebrain activations are reported at *p*<.005 uncorrected with k>59 voxels, equivalent to a FWE cluster correction for multiple comparisons of *p*<.05. The colorbar represents t-statistics. Inferior frontal cortex delineated in white with subregions delineated in black using the automated anatomical labelling (‘aal’) atlas. CG: control group; EG: experimental group; TMS: transcranial magnetic stimulation; IFC: inferior frontal cortex; IFGop: inferior frontal gyrus *pars opercularis*; IFGtri: inferior frontal gyrus *pars triangularis*; IFGorb: inferior frontal gyrus *pars orbitalis*; cTBS: continuous theta burst stimulation.

Indeed, the right IFC, more specifically the right IFGtri (see Methods for more details), was the global maximum when computing the aforementioned contrast in the CG (Fig.2**B-D**), and it was therefore used as a target region for the cTBS procedure of the EG instead of using an IFC region reported in the literature. The homologous IFC region in the left hemisphere (Fig.2**AC**) responded to a lesser extent to our contrast of interest and was therefore not targeted by the cTBS procedure. The cTBS procedure targeted the vertex [MNI *xyz* 0 −20 80] for the CG. Coil positioning is reported in Fig.2**D** for each group.

### Altered behavioral decisions on voice affective ambiguity after cTBS on the right IFC

We first assessed the effects of cTBS applied to the right IFC on the decisional patterns during the classification of neutral (no interest condition) as well as clear and ambiguous vocal affect. The behavioral classification data were parametrized along response speeds and decision probabilities in the three-alternative task (neutral, anger, or fear). Response speed and decision probability were analyzed using generalized linear mixed-effects (‘glmer’) and linear mixed-effects (‘lmer’) models, respectively. These models allowed us to test our hypothesis according to which participants of the CG would perform better and potentially faster at categorizing blended voices as opposed to those of the EG who received cTBS over the right IFC. As mentioned before, neutral voices were used as baseline control trials and we did not have any hypothesis concerning these, so we modelled these trials in all our analyses but discarded them from behavioral and neuroimaging data results. Their classification accuracy was very high (i.e., they were not misclassified as and mixed-up with the affective trials) and no significant difference between groups across runs was observed (χ^2^(1)=0.08, *p*=1; see Table 1). In the first statistical model with the probability of an anger choice on affective trials as the binomial dependent variable (Fig.1**B**), we observed significant effects along several of our experimental factors. First, we found significantly different choice probabilities between the morphing levels of the voice stimuli (factor *morphing*: χ^2^(5)=1496.55, *p*<.0001), with voices containing a higher proportion of anger being more likely classified as ‘anger’, while those containing a higher proportion of fear being more likely classified as ‘fear’ ([A_90_ > A_70_]: b=-0.11, χ^2^(1)=83.00, *p*<.0001; [A_90_ > AF_50_]: b=-0.25, χ^2^(1)=395.55, *p*<.0001; [A_90_ > F_70_]: b=-0.37, χ^2^(1)=870.09, *p*<.0001; [A_90_ > F_90_]: b=-0.42, χ^2^(1)=1075.91, *p*<.0001). These expected effects were observed independently of groups and runs. We then observed a significant main effect of the factor *run* (χ^2^(3)=8.56, *p*<.05) explained by the probability of an appropriate choice being higher for the base run as compared to the cTBS_post_ runs (b=0.76, χ^2^(1)=7.38, *p*<.001).

**Table 1.** Accuracy and reaction times data for neutral trials, for each group and run.

|  | **Base run** | **cTBS<sub>post</sub> Run 1** | **cTBS<sub>post</sub> Run 2** | **cTBS<sub>post</sub> Run 3** |
| --- | --- | --- | --- | --- |
| **CG acc** | 87.50% (6.55) | 87.78% (4.96) | 88.05% (5.55) | 88.05% (5.55) |
| **CG rt** | 1032ms (188) | 928ms (201) | 918ms (213) | 944ms (208) |
| **EG acc** | 93.23% (6.74) | 97.94% (5.10) | 97.06% (5.71) | 97.94% (5.10) |
| **EG rt** | 1112ms (192) | 1043ms (204) | 1061ms (186) | 1137ms (206) |

Value: mean (SD). acc: accuracy; rt: reaction times; ms: milliseconds.

We furthermore observed a significant *group* x *morphing* interaction (χ^2^(5)=67.05, *p*<.0001) explained by a better fit—namely, response probability matching the morphing percentage in each affective voice—of the choice probability of an anger choice for the EG compared to the CG for clear voices across all runs, namely A_90_ and F_90_ ([EG > CG] x [A_90_]: b=0.35, χ^2^(1)=3.86, *p*<.05; [EG > CG] x [F_90_]: b=-0.41, χ^2^(1)=5.54, *p*<.05) but not for the most ambiguous voices ([EG > CG] x [AF_50_]: b=0.06, χ^2^(1)=0.14, *p*>.10) or when comparing clear against ambiguous voices ([EG > CG] x [AF_50_ > A_90_, A_70_, F_90_, F_70_ > AF_50_]: b=-0.51, χ^2^(1)=1.07, *p*>.10).

Although the triple interaction for factors *group* x *morphing* x *run* was not significant (χ^2^(15)=13.60, *p*=.33), it however pertained to our main hypothesis of an impact of cTBS on ambiguous voice categorization. We therefore tested the contrast of a higher probability of an anger choice for the CG when categorizing most ambiguous as compared to least ambiguous voices—the inverse contrast yields to the same test with sign reversal of the difference—as a function of cTBS (contrast: [CG > EG] x [AF_50_ > A_90_, A_70_, F_90_, F_70_] x [cTBS_post_ > cTBS_pre_]) and found a significant effect (b=1.94, χ^2^(1)=5.24, *p*<.05). This result shows that there is a greater classification probability difference between clear and ambiguous voices for the EG as compared to the CG for cTBS_post_ vs. cTBS_pre_ runs.

Finally, we tested the effect of right IFC cTBS on choice probabilities specifically on the most ambiguous affective voices between groups, namely those containing fifty percent of fear and anger. This contrast confirmed that cTBS on the right IFC led to a difference between to classify the most ambiguous affective voices—with the CG showing a higher probability of classifying these as ‘anger’ voices as compared to the EG ([CG > EG] x [AF_50_] x [cTBS_post_ > cTBS_pre_]; b=0.31, χ^2^(1)=6.05, *p*<.05; see Fig.1**B**). In other words, this triple interaction contrast illustrates an improvement for the EG compared to the CG in classifying clear as opposed to ambiguous affective speech as a function of cTBS. The last contrast focused specifically on the most ambiguous voices and also shows a fundamental between-group difference triggered by cTBS on the right IFG with the CG’s probability of classifying AF_50_ voices as ‘anger’ being higher than that of the EG. Participants of the EG also improved their classification of the most ambiguous voices. This first model on the probability of an anger choice explained 15.85% of the variance including both fixed and random effects (R2c= 0.1585) and 10.23% of the variance solely for fixed effects (R2m= 0.1023).

Reaction time data were analyzed in a separate linear mixed-effects model with reaction times used as the dependent variable (Fig.3**C**), and we observed a main effect of *morphing* (χ^2^(5)=2126.79, *p*<.0001). This was explained by several pairwise differences between morphing levels (see Table S3) and generally illustrating a linear decrease of reaction times as a function of the increasing percentage of anger in the voice stimuli. The main effect of *run* was also significant (χ^2^(3)=58.54, *p*<.0001), showing overall slower reaction times for cTBS_pre_ than cTBS_post_ runs ([base run > cTBS_post_ run1]: b=550.07, χ^2^(1)=18.59, *p*<.0001; [base run > cTBS_post_ run2]: b=861.96, χ^2^(1)=45.48, *p*<.0001; [base run > cTBS_post_ run3]: b=833.80, χ^2^(1)=42.48, *p*<.0001). The *group* x *morphing* and *group* x *run* interactions were also significant (χ^2^(5)=40.67, *p*<.0001 and χ^2^(3)=8.27, *p*<.05, respectively). The former was explained by faster reaction times for the CG compared to the EG to categorize neutral voices (b=-537.23, χ^2^(1)=5.71, *p*<.05; all reaction times for neutral voices in Table 1), the latter by participants of the CG being faster than those of the EG to categorize voice stimuli of the cTBS_pre_ compared to cTBS_post_ run 1 (b=-296.61, χ^2^(1)=5.40, *p*<.05).

**Fig.3:**
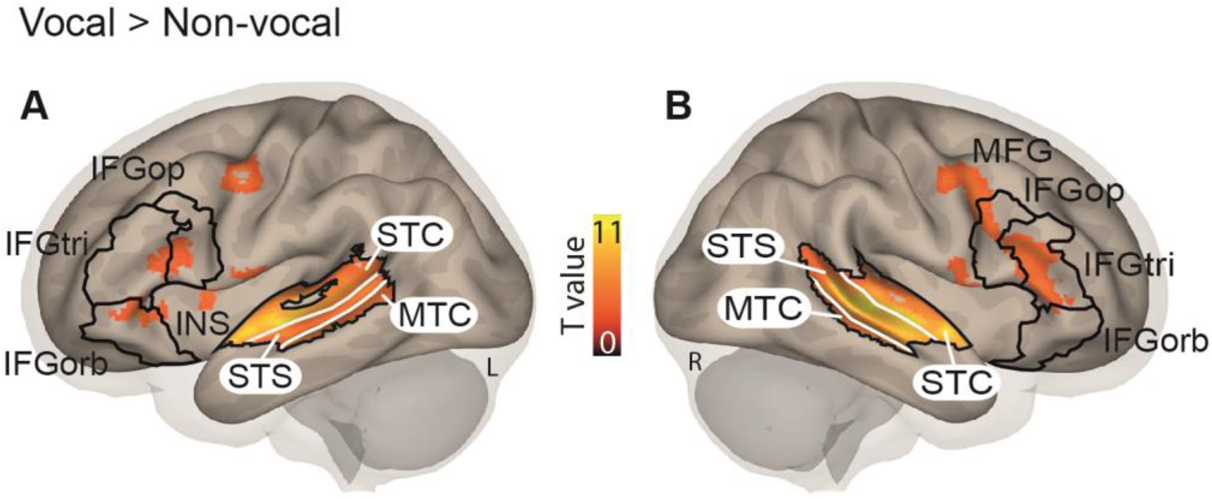
Sample-specific (N=35) activations for the VA localizer task. (**A**) Voice areas and IFC regions (black outline) in the left hemisphere for the [Vocal > Non-vocal] contrast. (**B**) Voice areas and IFC regions (black outline) in the right hemisphere for the [Vocal > Non-vocal] contrast. Wholebrain activations are reported at *p*<.005 uncorrected with k>59 voxels, equivalent to a FWE cluster correction for multiple comparisons of *p*<.05. Colorbars represent t-statistics. *IFGop* inferior frontal gyrus pars opercularis; *IFGtri* inferior frontal gyrus pars triangularis; *IFGorb* inferior frontal gyrus pars orbitalis; *INS* insula; *STC* superior temporal cortex; *STS* superior temporal sulcus; *MTC* middle temporal cortex; *DLPFC* dorsolateral prefrontal cortex; *MFG* middle frontal gyrus.

The triple interaction of interest was not significant (χ^2^(15)=10.98, *p*=.75), but since it was part of our main behavioral hypothesis, we computed the contrasts of interest using planned comparisons testing the effect of group, morphing, and run. Again, we ran the [CG > EG] x [AF_50_ > A_90_, A_70_, F_90_, F_70_] x [cTBS_post_ > cTBS_pre_] triple interaction contrast and the [CG > EG] x [AF_50_] x [cTBS_post_ > cTBS_pre_] contrast but none of them was significant (b=-440.48, χ^2^(1)=0.26, *p*>.10 and b=38.00, χ^2^(1)=0.09, *p*>.10, respectively). This second model explained 27.20% of the variance including both fixed and random effects (R2c=0.2720) and 10.12% of the variance solely for fixed effects (R2m=0.1012).

### Voice-sensitive activations in bilateral auditory cortex

To determine the regions in bilateral AC that are generally sensitive to voice compared to other types of auditory signals, we analyzed the data of a functional voice localizer scan specific to our sample. This voice localizer scan was specifically performed to spatially localize regions that might show specific increased activity for affective ambiguity expressed in voices in the main fMRI task. During this scan, participants listened to vocal and non-vocal sounds, and we found higher activity in bilateral STC and IFC (Fig.3**AB**; Table S4) when contrasting vocal against non-vocal sounds. This pattern of activations underlying the neural processing of voice sounds is similar to previously reported activation patterns and referred to as the “voice areas” (Belin, Zatorre et al. 2000) (VA). IFC subregions are also delineated in the maps of Fig.3.

### Functional brain activations for morphed voices classifications as a function of cTBS

We used the cortical definition of the VA to determine functional activations in the main experiment that were located inside as well as outside this general voice processing area. Imaging data were used to characterize brain mechanisms underlying the processing and classification of affective ambiguity using blended voices following cTBS over a specialized area, the right IFC.

Therefore, our contrast of interest was computed to uncover brain activity relating to [A_90_, A_70_, F_90_, F_70_ > AF_50_] x [cTBS_post_ > cTBS_pre_] and its opposite [AF_50_ > A_90_, A_70_, F_90_, F_70_] x [cTBS_post_ > cTBS_pre_], per group first and then between groups. We found enhanced brain activity in the former but not in the latter contrast. In fact, the [A_90_, A_70_, F_90_, F_70_ > AF_50_] x [cTBS_post_ > cTBS_pre_] contrast yielded enhanced brain activity in the STC (within the temporal voice areas), bilateral IFC especially in the *pars triangularis* for the CG (Fig.4**BE**) and EG (Fig.4**CF**), separately.

**Fig.4:**
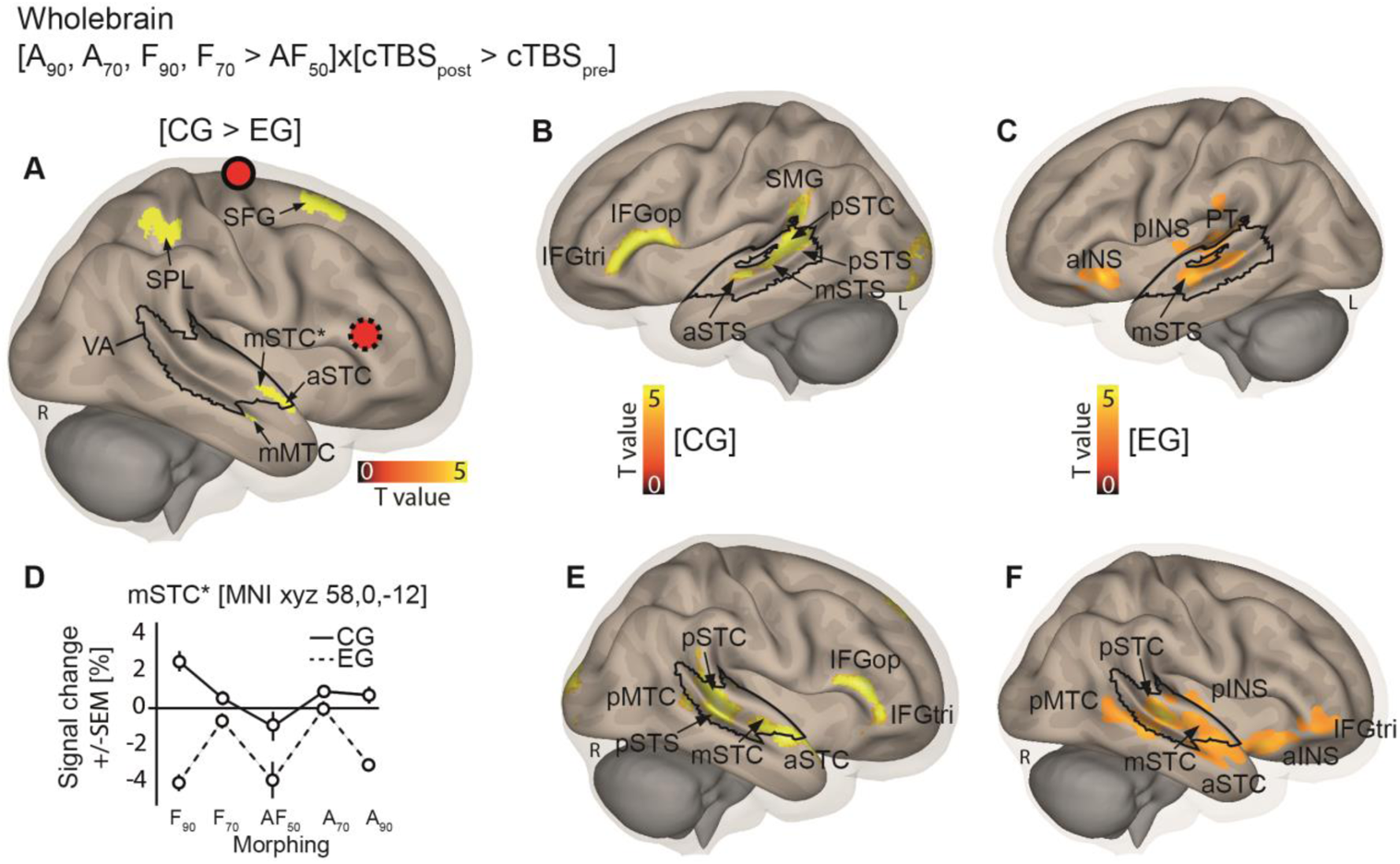
Neuroimaging results of the vocal affect classification task with outlines of sample-specific voice areas. (**A**) Whole-brain data showing between-group activations for the [CG > EG] x [A_90_, A_70_, F_90_, F_70_ > AF_50_] x [cTBS_post_ > cTBS_pre_] contrast with continuous theta burst stimulation (cTBS) over the vertex (red circle, black outline) for the control group (CG) as opposed to over the right inferior frontal gyrus (IFG; red circle, dotted outline) for the experimental group (EG). (**B**) Left hemisphere activations for [A_90_, A_70_, F_90_, F_70_ > AF_50_] x [cTBS_post_ > cTBS_pre_] contrast for the CG. (**C**) Left hemisphere activations for [A_90_, A_70_, F_90_, F_70_ > AF_50_] x [cTBS_post_ > cTBS_pre_] contrast for the EG. (**D**) Percentage of signal change (±SEM) extracted in the right mSTG* [MNI *xyz* 58, 0, −12] for the main contrast ([CG > EG] x [A_90_, A_70_, F_90_, F_70_ > AF_50_] x [cTBS_post_ > cTBS_pre_]). (**E**) Right hemisphere activations for the [CG > EG] x [A_90_, A_70_, F_90_, F_70_ > AF_50_] x [cTBS_post_ > cTBS_pre_] contrast for the control group. (**F**) Right hemisphere activations for the [CG > EG] x [A_90_, A_70_, F_90_, F_70_ > AF_50_] x [cTBS_post_ > cTBS_pre_] contrast for the experimental group. Sample-specific voice areas (VA) are outlined in each panel in black. *a*/*m*/*p* anterior/mid/posterior; *op*/*tri* pars opercularis/triangularis; *MFG* middle frontal gyrus; *STC* superior temporal cortex; *MTC* middle temporal cortex; *SPL* superior parietal lobule; *INS* insula; *PTe* planum temporale; *SMG* supramarginal gyrus; *STS* superior temporal sulcus. Wholebrain activations are reported at *p*<.005 uncorrected with k>59 voxels, equivalent to FWE cluster correction for multiple comparisons of *p*<.05. Colorbars represent t-statistics.

Computing the three-way interaction between the factors *group*, *morphing,* and *run* revealed activity within the VA (Fig.4**A**), located in anterior and mid STC as well as middle temporal cortex (MTC), and in superior parietal lobule and superior frontal gyrus only in the right hemisphere ([CG > EG] x [A_90_, A_70_, F_90_, F_70_ > AF_50_] x [cTBS_post_ > cTBS_pre_]; Fig.4**A**). The inverse contrast ([CG > EG] x [AF_50_ > A_90_, A_70_, F_90_, F_70_] x [cTBS_post_ > cTBS_pre_]) and the ([CG > EG] x [AF_50_] x [cTBS_post_ > cTBS_pre_] contrast did not yield any above-threshold brain activity. Peak coordinates and statistical information are reported in Table 2.

**Table 2.** Wholebrain activations for contrasts of interest, per group and between-groups.

| Region label | Hemisphere | MNI X | MNI Y | MNI Z | T-value | Cluster size (voxel count) |
| --- | --- | --- | --- | --- | --- | --- |
| SPL | R | 28 | -42 | 48 | 4.63 | 186 |
| STG | R | 58 | 0 | -12 | 3.74 | 84 |
| MFG | R | 20 | 20 | 56 | 3.73 | 73 |
| Cuneus | R | 18 | -80 | 32 | 3.53 | 120 |

| Region label | Hemisphere | MNI X | MNI Y | MNI Z | T-value | Cluster size (voxel count) |
| --- | --- | --- | --- | --- | --- | --- |
| STG | R | 58 | -26 | 2 | 5.70 | 549 |
| Cerebellum | L | -20 | -74 | -36 | 4.75 | 318 |
| Insula | R | 32 | 20 | -14 | 4.69 | 218 |
| STG | L | -50 | -16 | -6 | 4.49 | 686 |
| STS | R | 54 | -4 | -14 | 4.08 | 252 |
| AMY | R | 20 | -2 | -12 | 3.84 | 107 |
| IFGtri | R | 50 | 46 | -4 | 3.78 | 105 |

| Region label | Hemisphere | MNI X | MNI Y | MNI Z | T-value | Cluster size (voxel count) |
| --- | --- | --- | --- | --- | --- | --- |
| IFGtri | R | 54 | 34 | 10 | 5.21 | 324 |
| Cerebellum | L | -14 | 84 | -14 | 4.66 | 4273 |
| STS | R | 58 | -32 | 2 | 4.60 | 822 |
| STS | R | 54 | -4 | -12 | 4.38 | 372 |
| STG | L | -54 | -42 | 8 | 4.25 | 626 |
| IFGtri | L | -58 | 20 | 8 | 4.19 | 713 |
| STG | L | -50 | -20 | -6 | 3.82 | 66 |
| MFG | L | -4 | 64 | 28 | 3.47 | 98 |

Statistical threshold: $p < .05$ FWE cluster correction (voxelwise $p < .005$ uncorrected, $k > 59$ ).
*SPL* superior parietal lobule; *STG* superior temporal gyrus; *MFG* middle frontal gyrus; *STS* superior temporal sulcus; *AMY* amygdala; *IFGtri* inferior frontal gyrus *pars triangularis*; L: left hemisphere; R: right hemisphere.

### Functional connectivity data for morphed voice classifications as a function of cTBS

Functional connectivity analyses were computed in order to assess the *group* x *morphing* x *run* interaction on the organization of functional networks of ambiguous voice classification. Functional connectivity analyses allow an inference of the association—in the present case, positive or negative linear bivariate correlations with the task-dependent generalized psychophysiological interaction method—between regions observed in the wholebrain results. Seed regions of interest (ROI) overlapping with results from studies targeting the decoding of affective voices in the lateral, superior temporal cortex (Schirmer and Kotz 2006), medial temporal lobe (Frühholz, Trost et al. 2014) and more specifically the role of the IFC in vocal affect processing (Hoekert, Vingerhoets et al. 2010, Frühholz and Grandjean 2013) were selected (see Methods). We therefore ended up including eight ROI (bilateral IFC, bilateral anterior, mid and posterior STC, bilateral amygdala) in our seed-to-seed, generalized psychophysiological interaction analysis.

These analyses revealed anti-coupling in the right amygdala and left mid STC (Fig.5**AB**) as well as coupling in the left amygdala and right IFC (Fig.5**BC**) triggered by cTBS on the right IFC ([EG > CG] x [A_90_, A_70_, F_90_, F_70_ > AF_50_] x [cTBS_post_ > cTBS_pre_] contrast). This result indicated that when the EG as compared to the CG classified clear as opposed to ambiguous voices as a function of the cTBS procedure, linear negative correlation between left mid STC and right amygdala increased. On the other hand, linear association between the left amygdala and right IFC (the cTBS target region for the EG) increased. The cTBS procedure on the right IFC therefore seems to enhance functional connectivity between the bilateral amygdala and subparts of the VA and IFC. Since functional connectivity data were computed using bivariate correlations between neural nodes, inverse contrasts ([EG > CG] x [AF_50_ > A_90_, A_70_, F_90_, F_70_] x [cTBS_post_ > cTBS_pre_] or [CG > EG] x [A_90_, A_70_, F_90_, F_70_ > AF_50_] x [cTBS_post_ > cTBS_pre_]) yielded to a sign inversion (coupling in the right amygdala and left mid STC; anti-coupling in the left amygdala and right IFC) and were therefore not illustrated.

**Fig.5:**
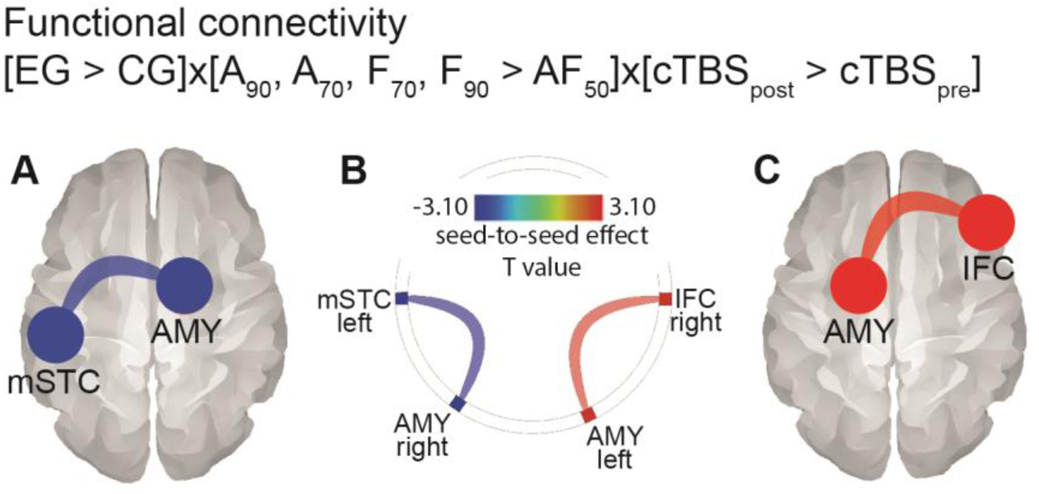
Seed-to-seed functional connectivity results for the vocal affect categorization task. (**A**) Anti-coupled functional connectivity was found between the left mid superior temporal cortex (mSTC) and the right amygdala (AMY) for the [EG > CG] x [A_90_, A_70_, F_90_, F_70_ > AF_50_] x [cTBS_post_ > cTBS_pre_] contrast. (**B**) Summary of coupled and anti-coupled functional connectivity data and statistical values. (**C**) Coupled functional connectivity between the left amygdala (AMY) and the right inferior frontal cortex (IFC) for the [EG > CG] x [A_90_, A_70_, F_90_, F_70_ > AF_50_] x [cTBS_post_ > cTBS_pre_] contrast. Colorbar shows two-tailed t-statistics. Statistical threshold: *p*<.05 FDR corrected for multiple comparisons at the seed (N=8) level.

## Discussion

The present study had the general aim of understanding the causal role of the inferior frontal cortex in classifying and representing speech at different levels of affective clearness and ambiguity. A combined cTBS-fMRI procedure was therefore used to alter activity in the right IFG *pars triangularis* for half of our participants (EG), while the other participants (CG) received cTBS over a control region, namely the vertex, which was supposed to not alter any neural activity relevant for affective sound processing and classification (Jung, Bungert et al. 2016). The cTBS procedure targeting the IFC led to an altered functioning of this region and affected decisional processes of the participants during the affect classification task. This target region was chosen because it was found active during the processing of affective voices in the CG, and it is a commonly found activation peak across many studies on voice processing, categorization, and classification (Schirmer and Kotz 2006, Frühholz and Grandjean 2012, Frühholz and Grandjean 2013, Suran, Rumiati et al. 2019, Grandjean 2020). The cTBS procedure had an impact on both behavioral decisions and cerebral processing. First, we found that cTBS on the right IFC improved voice classification, specifically for ambiguous affective voices. Second, IFC cTBS highlighted a reduced recruitment of STC regions for clear as opposed to ambiguous voices. Third, functional connectivity analyses for clear versus ambiguous voices revealed between-group differences, such that IFC cTBS led to anti-coupling between the mid STC and right amygdala and enhanced coupling between left amygdala and right IFC, the latter region being the target of the cTBS procedure in the EG.

The morphing procedure used to create affective ambiguity allowed us to assess if neural dynamics in IFC and connected regions would either represent affective voice classification challenges (as induced by ambiguous affective voices) or represent categorical affective certainty (as induced by clear affective voices). According to the first perspective, we hypothesized that IFC alteration by the cTBS procedure would potentially affect the classification of ambiguous affective voices, namely those with a blend of 50% anger and fear, which would represent the decisional difficulty of this condition. According to the second view, we hypothesized that altering IFC functioning using cTBS would affect the certainty of clear affective voice categorization, namely voices expressing mostly anger (90% anger and 10% fear voices) or mostly fear (90% fear and 10% anger voices). While these perspectives could be in opposition to each other, their mutual existence should also be considered in the IFC. In fact, fine-grained IFC mechanisms underlying the classification of ambiguous as opposed to clear affective voices could encompass both decisional difficulty and categorization certainty, respectively. This integrated view of the complementary roles and functions of the IFC in vocal emotion classification could also help explain research with unexpected categorization results—as well as the absence of expected results, for instance when altering left or right IFC using rTMS (Hoekert, Vingerhoets et al. 2010).

Given the abovementioned perspectives of the critical role(s) of the IFC as underlying behavioral decisions on and/or representations of affective voices, we first investigated the behavioral effects of cTBS on the right IFC, specifically the right IFGtri. We found a modulation of reaction times during the vocal affect decision task, but most importantly we found an influence on affect classification probabilities. These results highlight the direct impact of altered IFC functioning following cTBS, a region that seems on the one hand associated with the cognitive evaluation or judgement of affective voices (Frühholz and Grandjean 2012, Frühholz and Grandjean 2013, Swanborough, Staib et al. 2020) while the IFC is also known to be sensitive to nonverbal vocalizations in both children (Grossmann, Oberecker et al. 2010) and nonhuman primates (Romanski and Averbeck 2009), supporting sound classifications (Cohen, Hauser et al. 2006) and higher-order auditory representations (Cohen, Russ et al. 2009) on the other hand. Our reaction time data especially revealed that cTBS on the right IFC leads to faster reactions in the experimental run immediately after the application of cTBS for the EG. The neural effects of cTBS are usually largest immediately after the application, and the effects are known to decay over time—they can last up to 60 minutes after this specific type of patterned stimulation (Huang, Edwards et al. 2005). If the IFC is assumed to be a neural node for lifting affective voice processing on a cognitive-focused level (Ethofer, Anders et al. 2006, Schirmer and Kotz 2006, Brück, Kreifelts et al. 2011, Frühholz and Grandjean 2013) that demands processing efforts (Verbruggen, Aron et al. 2010)—as observed in spoken word recognition and phonetic competition (Zhuang, Tyler et al. 2012, Rogers and Davis 2017, Xie and Myers 2018), inhibition of the IFC with cTBS might loosen this cognitive focus to facilitate more intuitive processing (Zander, Horr et al. 2016) with presumably better processing efficiency (Dippel and Beste 2015). Our classification choice probability data might also point in a similar direction. We especially observed a better classification probability for the clearest affective voices after cTBS in the EG group compared to the CG. After cTBS, affective decisions of participants much more follow the quantity of sensory acoustic evidence. This might indicate that these decisions are more sensory driven rather than driven by high-level cognitive processing. This also indicates that the IFC normally seems indeed a neural node for high-level cognitive evaluation of sound information and affective values of voice signals but seems to flag the outcome of such evaluation with more categorical uncertainty, especially for more complex affective categorization tasks. These observations together would suggest a mixed functional role of the IFC according to the two primary hypotheses on the IFC: the IFC would both compute cognitive evaluation on socio-affective voice information, but it would also tag such information with flags of affective uncertainty and decisional precaution.

The relevance of this IFC subregion as targeted by our cTBS procedure for complex affective evaluation and classification tasks is highlighted by previous studies. The IFC is a rather large region (Frühholz and Grandjean 2013) and it includes several subregions or subparts such as its more superior part, *pars opercularis*, the more inferior and anterior part, *pars triangularis* and the most ventral part, *pars orbitalis* that is located next to the orbitofrontal cortex. In our study, the location of the IFC was determined *a priori* to serve as cTBS target region and it was exclusively within the *pars triangularis* in the right hemisphere. In another study, this specific subregion of the right IFC was recruited when more complex affective voice categorizations as opposed to simpler discrimination was performed, also sometimes labelled ‘unbiased’ versus ‘biased’ perceptual decision making, respectively (Dricu, Ceravolo et al. 2017, Gruber, Debracque et al. 2020). Such results are therefore in line with our data, especially when considering the fact that an affect classification task was employed in our procedure. Affective voice discrimination as opposed to categorization would on the other hand depend on a different IFG subregions, namely the bilateral IFG *pars opercularis* (Dricu, Ceravolo et al. 2017), to which only residual TMS current could be transmitted due to the cTBS procedure and coil location in our study (see Methods). According to our results and to the literature (Frühholz and Grandjean 2013, Dricu, Ceravolo et al. 2017, Frühholz and Schweinberger 2020, Gruber, Debracque et al. 2020), the right and potentially the left IFG *pars triangularis* (Lupyan, Mirman et al. 2012) could therefore be highly selective to categorization, by flagging the outcome of cognitive-driven affective evaluation processes with some level of uncertainty (Toelch, Bach et al. 2013, Nastase, Iacovella et al. 2014).

In a more general sense, the IFC was previously also causally linked to social perception and affective processing in social groups. In fact, repetitive TMS over the left IFC revealed perturbed social perception while social cognition was preserved, with slower reaction times observed for emotion recognition (Keuken, Hardie et al. 2011). A similar TMS procedure targeting the *pars opercularis* of the left IFG led to faster categorization of negative social groups and disrupted semantic priming for negative words, as opposed to TMS over the vertex (Suran, Rumiati et al. 2019). These two studies emphasize the importance of the target location for a TMS procedure—and add nuance to IFC subparts functioning and specificity—by pointing toward slower or faster responses according precise IFC disruption in the *pars orbitalis* (Keuken, Hardie et al. 2011) or *opercularis* (Suran, Rumiati et al. 2019), respectively.

Considering our neuroimaging data and when looking at groups separately (CG, EG), bilateral IFG *pars triangularis* activity was observed specifically for clear as compared to ambiguous voices in both groups separately, but not in the inverse contrast. This observation seems contrary to the hypothesis of the IFC being pre-dominantly a brain node coding and regulating decisional challenges especially during sensory ambiguity. The IFC is active when making socio-affective decisions on voice signals, but not simple acoustic decisions (Roswandowitz, Swanborough et al. 2020), which would again point to the notion of the IFC of representing affective categorical information of voice signals, but maybe flagged with categorical uncertainty and choice precaution (Toelch, Bach et al. 2013, Nastase, Iacovella et al. 2014). This might concern especially the right IFC, but some bilateral interaction of homologue areas in left and right IFC could be expected, as we revealed lower bilateral IFC activity in the EG group. The bilateral functional significance of the IFC could be influenced solely by right hemisphere cortical interference through cTBS, as already shown using TMS on the left prefrontal cortex (Nahas, Lomarev et al. 2001). Such assumptions should however be tested in the future either by including a group of participants undergoing cTBS over the left IFC or by using intermittent TBS, which is known to enhance brain activity and not alter it (Annika, Alia et al. 2010, Oberman, Edwards et al. 2011), over the right and/or left IFG *pars triangularis*. Such assumption would however go against inter-hemispheric plasticity, as observed in speech production with right IFC recruitment following left IFC alteration (Hartwigsen, Saur et al. 2013), but it seems that the impact of stimulation intensity could have a crucial impact on current transmission in the brain tissues (Nahas, Lomarev et al. 2001).

We designed an experimental task that would specifically test the causal role of the IFC in classifying ambiguous and/or clear affective speech and flagging the decision with some weighting of uncertainty, but numerous other functions of this brain region have been proposed and some of them could apply to our data (Liakakis, Nickel et al. 2011). For instance, behavioral classifications imply not only categorical associations but also action and motor functioning as well as executive functions, most notably inhibition. Inhibition was shown to causally rely on the right IFC, with direct current stimulation over the IFC causing an ‘activation of inhibition’ (Jacobson, Javitt et al. 2011). Such topic was also reviewed in detail and while the distinct role of prefrontal cortex subregions was initially questioned (Aron, Robbins et al. 2004), the specific role of the IFC would be to implement some cautionary mechanisms over immediate response tendencies (Aron, Robbins et al. 2014). According to this literature and to our data, it is therefore possible that cTBS over the right IFG *pars triangularis* enabled the release of inhibition in the EG. Response inhibition and inhibition of immediate categorical decisions might have appeared in the CG, leading to more conservative decisions and a more cognitive-driven processing especially concerning clear affective voices. In the EG, cTBS might have led to a release of inhibition and maybe to a switch from a cognitive to a more intuition-based processing of affective speech. These two processing modes of a cognitive-driven and intuition-driven processing of sensory information have been documented in the literature (Gilovich, Griffin et al. 2002, Kahneman 2002) and are potentially on a temporal continuum (Hogarth 2010), with intuition-driven processing supposed to be more efficient (Hammond, Hamm et al. 1987), especially when relying on prior expertise (Dane, Rockmann et al. 2012). A higher processing efficiency of clear affective speech in our study might be indicated by a significantly lower activity in anterior STC in the EG. The anterior STC is a major brain node in the analysis of affective speech (Ethofer, Bretscher et al. 2012, Frühholz, Trost et al. 2016), and shows functional (Ethofer, Bretscher et al. 2012) and structural (Ethofer, Bretscher et al. 2013, Frühholz, Gschwind et al. 2015) connections to the right IFC. Especially, the between-group contrasts highlighted above-threshold voxels in the right mid-to-anterior STC for CG compared to EG, especially when comparing the cTBS-dependent classification of clear as opposed to ambiguous voices—but not for the inverse contrast. The role of the STC in affective speech decoding is well documented (Grandjean, Sander et al. 2005, Schirmer and Kotz 2006, Ethofer, Bretscher et al. 2012, Frühholz, Ceravolo et al. 2012, Witteman, Van Heuven et al. 2012, Ceravolo, Frühholz et al. 2016, Frühholz and Schweinberger 2020, Grandjean 2020) and among this vocal emotion literature, some studies isolated activity enhancement in the right anterior STC for the categorization of female-ambiguous voices by male participants (Sokhi, Hunter et al. 2005). Right anterior STC activity was also enhanced as a function of trial-level acoustic distance between voices when categorizing morphed male and female voices to create gender-ambiguous stimuli (Charest, Pernet et al. 2013). Perceived gender ambiguity did however recruit the bilateral IFG and the posterior and anterior cingulate cortex. These results were interpreted by the authors as a two-stage process for gender voice classification, with auditory feature extraction in the anterior STC and voice categorization in the bilateral IFG, in a similar fashion described by Schirmer and Kotz for processing and making decisions on affective speech (Schirmer and Kotz 2006). While this interpretation was focused on categorizing gender and gender-ambiguity, it is well suited to our results even though our morphing procedure targets vocal emotion. It also means that mid and anterior STC neurons have the ability to influence classification and decision processes in the IFC and that at some point altered functioning in the IFC—as caused by cTBS in our study for the EG—could be overcome or at least reduced by right anterior STC activity and its feature-selectivity function.

Our functional connectivity results also highlight the importance of the limbic system for processing affective speech. The right amygdala showed decreased functional connectivity with the left STC in the EG vs. CG for the classification of clear and ambiguous voices, while the left amygdala was significantly anti-coupled with the right IFC. These results are coherent with existing literature on the brain networks of vocal affect processing (Frühholz, Ceravolo et al. 2012, Frühholz, Trost et al. 2014) and argue for a wider role of the limbic system in affective voice classification (Frühholz, Trost et al. 2014) and feature extraction (Pannese, Grandjean et al. 2016) depending on sensory ambiguity of the voice signal. In fact, automatic, non-voluntary, and presumably intuitive processing of vocal affect, especially anger (Grandjean, Sander et al. 2005, Sander, Grandjean et al. 2005), would take place in the amygdala following which feature extraction would be completed in the STC (Schirmer and Kotz 2006). According to Schirmer and Kotz (Schirmer and Kotz 2006), the last of the three-stage process of vocal affect processing would consist of an evaluative process in the IFC, allowing the participants to make a decision. For the categorization of affective speech ambiguity, our results show that such final stage is enhanced by temporary interference in the right IFC with coupling occurring between the left amygdala and right IFC in the EG compared to the CG. This observation suggests a potential enhancement in feedback generation from automatic processing—in the amygdala—and final decision leading to categorization, taking place in the right IFC.

In conclusion, our data might provide a more detailed and presumably mechanistic picture about the functional role of the IFC in social sound cognition and especially in voice signal classification along socio-affective dimensions. While previous research has suggested partly opposing roles of the IFC either for coding decisional challenges for difficult sound classifications or for high-level sound and voice categorical representations, the mechanistic functional role of the IFC might fall in-between these two perspectives. When applying cTBS to the right IFC in humans, their decisional responses on clear and ambiguous affective speech become faster immediately after cTBS, they follow more closely sensory evidence especially for clear voices, they show significantly lower activity in the auditory cortex, and they show higher fronto-limbic connectivity. Taken together, these data suggest that inhibition of the IFC leads to a more intuitive—and potentially more efficient—processing mode in the EG compared to a more cognitive- and analysis-driven processing in the CG. In terms of an involvement of the IFC in normal affective speech processing, the IFC might implement a cognitive processing mode that represents affective categories with more caution, and this categorical and representational caution might be shared with the anterior STC and the amygdala as major nodes in the neural affective speech analysis machinery.

## Material and Methods

### Participants

Forty healthy volunteers took part in the fMRI study. It was shown that 24 participants for an fMRI study accounted for at least 80% of the variance at the voxel level, as estimated for a conservative [0.01<α<0.05] alpha (Desmond and Glover 2002). Sample size was therefore calculated using G*Power 3 software (Faul, Erdfelder et al. 2007) with standard values for the architecture of our study (power of 0.95 and alpha of 0.05) and 80% of explained variance was reached with a sample of 20 participants per group.

We hence recruited 40 participants matching our inclusion criteria. Half of them were randomly assigned to the control group (CG) while the remaining participants were assigned to the experimental group (EG). For each group, 10 female and 10 male participants were included. Five participants were excluded from the final sample due to corrupted log files (N=2) and the impossibility to determine a reliable motor threshold (N=3). Therefore, the final groups included 18 participants for the CG (mean age=22.35, SD=1.66, 8 female) and 17 participants for the EG (mean age=25.05, SD=5.18, 7 female) with no significant between-group age difference (F(1,34)=3.66, *p*=.065). Participants were informed of all aspects of the experiment before giving their informed written consent to take part in the study. They were also informed that they could abort the experiment and quit the study at any time without justification. The study was conducted according to the Declaration of Helsinki and approved by the Ethics Committee of the University of Zürich, Switzerland.

### Vocal affect categorization task

Our stimuli consisted of voice utterances (“Aah”) from the Montreal Affective Voices database (Belin, Fillion-Bilodeau et al. 2008). Voices were pronounced by five female and five male actors. They expressed either a neutral emotional tone or a mix of anger and fear with a varying percentage of each emotion (percentage of anger/fear): 90/10, 70/30, 50/50, 30/70, 10/90 (Fig.1), respectively labelled A_90_, A_70_, AF_50_, F_70_, F_90_ in the text. Each voice was morphed using the same actor to avoid creating a strange identity that would add a critical confound to the emotional morphing procedure.

For each run of the vocal affect categorization task, we therefore had a total of five conditions of interest involving a morphing of anger and fear emotions (24 trials each) and one control condition involving neutral voices only (10 trials). Each voice had a duration of 500 ms to 1200 ms, the order of which being pseudo-randomized for each run and participant (Fig.1**A**). Each run included 24 trials of each conditions of interest for a grand total of 72 trials and 30 neutral voice trials across the three experimental runs and the control run.

The experiment took place at the University Hospital of Zürich, Switzerland, using the research-dedicated whole-body magnetic resonance imaging (MRI) scanner (Philips Achieva 3T) of the Laboratory for Social and Neural Systems Research (Department of Economics, University of Zürich, Zürich, Switzerland). Participants were first taken to a transcranial magnetic stimulation (TMS) room in order to determine their individual motor threshold through a standard procedure (see TMS procedure below). They were then taken to the MRI room and comfortably installed in the scanner. The MRI session started with one base run, followed by the TMS procedure and the three experimental runs (Fig.1**A**). At the end of the control run, the participants stayed on the scanner table. The table was taken halfway out and the TMS apparatus was brought to the participant. For the CG, continuous theta burst stimulation (cTBS) was administered for 40 seconds over the vertex (MNI *xyz* [0, −20, 80]) because it was previously successfully used as a good control site for TMS protocols(Jung, Bungert et al. 2016). For the EG, cTBS was administered to the right inferior frontal cortex (rIFC; MNI *xyz* [54, 34, 10]), for 40 seconds as well (see detailed TMS procedure below). As soon as the TMS procedure ended (for both groups), necessary care was taken to relocate the participant as fast and as smoothly as possible inside the scanner magnet. This procedure was used to minimize the time between stimulation and the start of the first experimental run, in order to have the most reliable TMS effect over the rIFC for the following runs across participants. The average duration between the end of the TMS procedure and the start of the first experimental run was 3.33 minutes (SD=0.47) for the CG and 3.00 minutes (SD=0.52) for the EG. No statistical difference was observed regarding this duration between groups (F(1,34)=0.33, *p*=.57).

For all runs, the task of the participants was to explicitly categorize the emotional tone of the voice that was presented in each trial by a key press using three buttons on an MRI-compatible response box (Current Designs Inc., PA, USA). Button mapping was fully randomized between participants. Buttons represented the following options: Anger, Fear, Neutral. Participants were instructed to respond as fast and as accurately as possible during the 2 sec blank screen that appeared after each trial (see Fig.1**A**).

### Behavioral data analysis

To take into account within- and between-subject variance for the initial stimulus evaluation and for the main cTBS-fMRI task—for the dependent variable of each model—we used lmerTest (Kuznetsova, Brockhoff et al.) and lme4 packages (Bates, Maechler et al. 2014) in R Studio (Team 2014) to perform mixed-effects modelling of the data. For all analyses, models were tested using type II Wald Chi-square tests using the ‘Anova’ function of the ‘car’ package (Fox, Friendly et al. 2007). We report effect sizes using the ‘MuMIn’ (Barton and Barton 2015) package based on two indicators, a marginal and a conditional R2 (R2m and R2c, respectively). R2m reflects the variance explained by the fixed factors, R2c the variance explained by the entire model (both fixed and random effects).

#### Initial stimulus evaluation and selection

For the initial evaluation of the stimuli including all possible morphing levels in steps of 10% as evaluated by a sample of nineteen participants, we used a generalized linear mixed-effects model (‘glmer’) for voice categorization. In this model, the dependent variable was the raw emotion categorization response of our sample of participants, with ‘1’ coding an ‘anger’ response and ‘2’ a ‘fear’ response. Fixed effects included the Morphing factor with eleven morphing levels of the voice stimuli (anger/fear percentage: 100/0, 90/10, 80/20, 70/30, 60/40, 50/50, 40/60, 30/70, 20/80, 10/90, 0/100) with N=10 trials per morphing level (N=110 trials per participant). Random effects included in this order: Participant, Age, Sex, Stimulus identity, Stimulus speaker sex. The same model was used to analyze reaction time data—these were of no interest for stimulus selection—with reaction times as dependent variable. All reported *p*-values of these analyses use a Bonferroni correction for multiple comparisons implemented in package ‘lmerTest’.

#### Vocal affect categorization task

For the main cTBS-fMRI task, mixed-effects modelling of both the responses (model 1, binomial modelling, generalized linear mixed-effects model ‘glmer’) and reaction times (model 2, linear modelling, linear mixed-effects model ‘lmer’) of our participants were computed, for each trial. Data included our conditions of interest, namely the 90/10, 70/30, 50/50, 30/70, 10/90 anger/fear morphed voices—labelled A_90_, A_70_, AF_50_, F_70_, F_90_, respectively, in the manuscript— and the control voices expressing neutral content (no morphing, filler condition of no-interest). For reaction times data and since raw values were not normally distributed, the log of the reaction times was used. For model 1, raw responses were used to determine the accuracy for an anger response for each trial, in a binomial manner. This binomial variable—coded ‘1’ for correct and ‘0’ for incorrect responses—was the dependent variable while the log of the reaction times was the dependent variable for model 2. For both models, fixed effects included, in this order, the interaction between Group, Morphing and Run while Participant, Age and Sex were introduced as random effects. These models allowed us to test our hypothesis according to which participants of the CG would perform significantly better at categorizing morphed voices, taking into account the TMS procedure: [CG > EG] x [AF_50_ > A_90_, A_70_, F_90_, F_70_] x [cTBS_post_ > cTBS_pre_] and its inverse regarding morphing [CG > EG] x [A_90_, A_70_, F_90_, F_70_ > AF_50_] x [cTBS_post_ > cTBS_pre_]. For these two contrasts, the weighting of the vector for the Morphing factor was similar to the one used for fMRI data, namely [6*(AF_50_) > −2*(A_90_), −1*(A_70_), −2*(F_90_), −1*(F_70_)] and [-6*(AF_50_) > 2*(A_90_), 1*(A_70_), 2*(F_90_), 1*(F_70_)]. A contrast targeting specifically the most ambiguous voices: [CG > EG] x [AF_50_] x [cTBS_post_ > cTBS_pre_].

All reported *p*-values of these behavioral analyses use a Bonferroni correction for multiple comparisons implemented in package ‘lmerTest’.

### Voice-sensitive areas localizer task

#### Stimuli

Auditory stimuli consisted of sounds from a variety of sources. Vocal stimuli were obtained from 47 speakers: 7 babies, 12 adults, 23 children and 5 older adults. Stimuli included 20 runs of vocal sounds and 20 runs of non-vocal sounds. Vocal stimuli within a run could be either speech 33%: words, non-words, foreign language or non-speech 67%: laughs, sighs, various onomatopoeia. Non-vocal stimuli consisted of natural sounds 14%: wind, streams, animals 29%: cries, gallops, the human environment 37%: cars, telephones, airplanes or musical instruments 20%: bells, harp, instrumental orchestra. The paradigm, design and stimuli were obtained through the Voice Neurocognition Laboratory website (http://vnl.psy.gla.ac.uk/resources.php). Stimuli were presented at an intensity that was kept constant throughout the experiment 70 dB sound-pressure level.

#### Experimental procedure, paradigm

Participants were instructed to actively listen to the sounds, both vocal and non-vocal. The distinction between vocal and non-vocal runs was not revealed to the participants who were therefore naïve to run organization and timing. The silent inter-run interval between each run of either vocal or non-vocal was 8s long. Task duration was about 10 minutes in total. Wholebrain, sample-specific result outline of this task for the vocal > non-vocal contrast of interest are reported in Fig.3 and voice areas are outlined in black in Fig.4.

### TMS procedure

As aforementioned, participants were stimulated either over the rIFC (EG) or over the vertex (CG) by means of standard cTBS (Huang, Edwards et al. 2005). First, we determined the active motor threshold (aMT) individually for each participant by stimulating the primary motor cortex in the left hemisphere. The aMT was defined as the percent of maximum stimulator output (mean intensity was 49.3%, SD 6.52%) required to elicit a motor-evoked potential larger than 200μV from the contralateral first dorsal interosseous muscle in five out of ten TMS pulses. During the determination of the aMT the participants exerted a constant pressure between the index finger and the thumb of about 20% of the maximum force (Huang, Edwards et al. 2005). For the cTBS protocol in the MRI room, the stimulation intensity was set to 80% of the aMT (mean intensity was 39.94%, SD 5.21%). Due to unwanted muscle contraction and discomfort at the right IFGtri location, percentage of aMT for the EG was further reduced by 10%, leading to lower amplitude stimulation for the EG compared to the CG in the MRI scanner (CG: mean intensity=42.84, SD=4.78; EG: mean intensity=36.87, SD=3.75; F(1,34)=16.80, *p*<.001).

On the bed of the MRI scanner, the participants received cTBS with an MR-compatible coil (MRi-B91 coil, MagVenture A/S, Farum, Denmark). Stimulation site and coil orientation (see TMS site localization; Fig.2) were marked on a fixed cap by means of a TMS Neuronavigation system (BrainSight 2, Rogue Research Inc., Canada). The cTBS stimulation protocol comprised bursts of 3 stimuli at 50Hz that were repeated with a frequency of 5Hz for 40s, resulting in a total of 600 pulses. The implemented cTBS protocol is thought to reduce the excitability of the stimulated brain region for about 60min (Huang, Edwards et al. 2005).

### TMS site localization

We determined the stimulation sites using individual T1-weighted structural scans and TMS Neuronavigation system (BrainSight 2, Rogue Research Inc., Canada). We used data from the CG—acquired before those of the EG—to define the coordinates of the rIFC (MNI *xyz* 54 34 10), coordinates that also corresponded to those observed in a previous study on the role of the rIFC for emotional judgments (Hoekert, Vingerhoets et al. 2010) and this location also overlapped with vocal judgments in general (Schirmer and Kotz 2006) and vocal emotion processing in the inferior frontal gyrus (Frühholz and Grandjean 2013). For each participant, we transformed the rIFC peak coordinates into the native space of the individual structural scan using the parameter estimates for spatial normalization of the anatomical scan performed in SPM12. The TMS coil was positioned tangentially to the cortical surface over the rIFG, with the handle perpendicular to the rIFC. As a control site we used the vertex, which was defined as the meeting point of the pre- and post-central sulcus in the interhemispheric fissure. For the control group, the TMS coil was positioned tangentially to the cortical surface over vertex (MNI *xyz* [0, −20, 80]), with the handle pointing in a posterior direction (see Fig.2**D**).

### MRI data acquisition

Imaging data acquisition was performed at the Laboratory for Social and Neural Systems research of the University of Zürich, on a Philips Achieva 3T whole-body scanner equipped with an eight channel MR head coil. Four runs of 10 min each were collected for each participant (1 base run, 3 experimental runs). Each run contained 200 volumes (voxel size = 3 x 3 x 3mm3, 0.5mm gap, matrix size = 80 x 80, TR/TE = 2100/30ms, flip angle = 79, parallel imaging factor = 1.5, 35 slices acquired in ascending order for full coverage of the brain). High-resolution T1-weighted 3D turbo field echo structural scans were acquired and used for image registration and normalization (181 sagittal slices, matrix size = 256 x 256, voxel size = 1mm3, TR/TE/TI = 8.3/2.26/181ms).

### MRI data analysis

Functional images were analyzed with Statistical Parametric Mapping software (SPM12, Wellcome Trust Centre for Neuroimaging, London, UK, http://www.fil.ion.ucl.ac.uk/spm). Preprocessing steps included realignment to the first volume of the time series, slice timing, normalization to the Montreal Neurological Institute (MNI; Collins, Neelin et al.) space using the DARTEL toolbox (Ashburner 2007) and spatial smoothing with an isotropic Gaussian filter of 8mm full width at half maximum. To remove low frequency components, we used a high-pass filter with a cutoff frequency of 128s. Anatomical locations were defined with a standardized MNI coordinate database using xjView toolbox (https://www.nitrc.org/projects/xjview).

#### Voice-sensitive areas localizer task

For the voice-sensitive areas localizer task, a general linear model was used to compute first-level statistics, in which each run was modelled by using a run function and was convolved with the hemodynamic response function, time-locked to the onset of each run. Separate regressors were created for each condition (vocal and non-vocal; Condition factor). Finally, six motion parameters were included as regressors of no interest to account for movement in the data. The Condition regressors were used to compute simple contrasts for each participant, leading to a main effect of vocal and non-vocal material at the first-level of analysis [1 0] for vocal, [0 1] for non-vocal. These simple contrasts were then taken to a flexible factorial second-level analysis in which there were two factors: the Participants factor with independence set to yes, variance set to unequal and the Condition factor with independence set to no, variance set to unequal.

#### Vocal affect categorization task

For the experimental runs and the base run of the main task, we used a first-level general linear model, in which each stimulus display was modelled by using a stick function and was convolved with the hemodynamic response function. Events were time-locked to the onset of the voice stimuli. Separate regressors were created for each condition of interest (five conditions with percentage of anger/percentage of fear: 90/10, 70/30, 50/50, 30/70, 10/90 or respectively A_90_, A_70_, AF_50_, F_70_, F_90_) and for neutral voices (regressor of no-interest). We thus had five regressors of interest including 24 trials each per run (base and experimental runs, respectively cTBS_pre_ and cTBS_post_ runs) and 72 trials each in total for the 3 experimental runs, in addition to the neutral voice condition as non-interest regressors (10 trials per run, 40 trials in total). Moreover, six motion parameters were included as regressors of no interest to account for movement in the data. Our design matrix was therefore as follows: Anger/Fear 90/10 (A_90_), Anger/Fear 70/30 (A_70_), Anger/Fear 50/50 (AF_50_), Anger/Fear 30/70 (F_70_), Anger/Fear 10/90 (F_90_), Neutral, Movement parameters, Constant term; 11 columns in total per run. We therefore had four sessions per model (base run, experimental run 1, 2, 3), leading to 44 columns in the design matrix. Each regressor of interest (Anger/Fear conditions) was used to compute contrasts for each participant (first-level statistics). First-level contrasts were computed to highlight the difference between difficult (more ambiguous) and easier (less ambiguous) trials for the emotional categorization of voices in all runs, especially the experimental runs. This procedure was decided based on the fact that our TMS inhibitory effect of the rIFC would last approximately 60 min (Huang, Edwards et al. 2005) and our runs were 10 min each for a total of 30 min, thus clearly within the bounds of the TMS effect. The contrast of interest was therefore the following: [A_50_ > A_90_, A_70_, F_90_, F_70_] x [cTBS_post_ > cTBS_pre_] (contrast vector: −2 −1 6 −1 −2 for each experimental run) and we computed its inverse regarding morphing, [A_90_, A_70_, F_90_, F_70_ > AF_50_] x [cTBS_post_ > cTBS_pre_] (contrast vector: 2 1 −6 1 2 for each experimental run) for each group separately as well as a vector specifically for the most ambiguous voices ([AF_50_] x [cTBS_post_ > cTBS_pre_], contrast vector: 0 0 1 0 0 for each experimental run). The weighted contrasts were used to better characterize the level of morphing of the voice stimuli as a function of the BOLD signal.

First-level contrast results of each participant were then averaged by group at the second-level using a two-sample t-test analysis. Using this procedure allowed us to test for an effect of cTBS stimulation over the vertex (CG) as opposed to cTBS over the rIFC (EG), our region of interest thought to be responsible for sensitive, accurate vocal emotional judgments. We therefore looked at the interaction between our contrasts of interest between groups: [CG > EG] x [AF_50_ > A_90_, A_70_, F_90_, F_70_] x [cTBS_post_ > cTBS_pre_] (contrast vector: 1 −1), [CG > EG] x [A_90_, A_70_, F_90_, F_70_ > AF_50_] x [cTBS_post_ > cTBS_pre_] (contrast vector: −1 1), [CG > EG] x [AF_50_] x [cTBS_post_ > cTBS_pre_] (contrast vector: 1 −1). Second-level statistical analyses of the main task assumed that Participants (Factor 1) were independent whereas Conditions (Factor 2) were not. Variance estimation was set to unequal for all factors in order to take into account the inhomogeneous variance of the data.

For the voice-sensitive areas localizer task, simple contrasts were then taken to a flexible factorial second-level analysis in which there were two factors: the Participant factor with independence set to yes, variance set to unequal and the Voice factor with independence set to no, variance set to unequal.

All wholebrain activations are reported at a threshold of *p*<.005 (uncorrected) and a cluster extent threshold of k > 59 voxels, equivalent to a Family-Wise Error correction for multiple comparison of *p*<.05 at the cluster level. This threshold was based on the final FWHM of the data (11.9, 11.9, 11.1 mm), using the ‘3dClustSim’ function in AFNI (http://afni.nimh.nih.gov/afni) software (Cox 1996), using a non-parametric method with 10’000 iterations to estimate the necessary cluster extent thresholding for side-to-side voxels (NN-2 option). ‘3dClustSim’ reports a cluster extent threshold for each specified statistical *p*-value and follows the assumption that neighboring voxels are part of a similar functional response pattern, rather than a completely different and independent measure as implied by the family-wise error correction at the voxel level. Inferior frontal cortex delineated (Fig.2) using the automated anatomical labelling (‘aal’) atlas (Tzourio-Mazoyer, Landeau et al. 2002).

### Functional connectivity analysis

Seed-to-seed functional analysis was performed for all runs (base and experimental) using the CONN toolbox (Whitfield-Gabrieli and Nieto-Castanon 2012) version 19.b implemented in Matlab 9.0 (The MathWorks, Inc., Natick, MA, USA). Functional connectivity analyses were computed using as seeds each region of interest (ROI) overlapping with results from studies targeting the decoding of emotional prosody in the lateral, superior temporal cortex (Schirmer and Kotz 2006), medial temporal lobe (Frühholz, Trost et al. 2014) and more specifically the role of the IFC in vocal emotion processing (Hoekert, Vingerhoets et al. 2010, Frühholz and Grandjean 2013). We therefore ended up including eight ROI (bilateral IFC, bilateral anterior, mid and posterior STC, bilateral amygdala) in our seed-to-seed, generalized psychophysiological interaction analysis. Spurious sources of noise were estimated and removed using the automated toolbox preprocessing algorithm, and the residual BOLD time-series was band-pass filtered using a low frequency window (0.008 < f < 0.09 Hz). Correlation maps were then created for each condition of interest by taking the residual BOLD time-course for each condition from atlas regions of interest and computing bivariate Pearson’s correlation coefficients between the time courses of each voxel of each ROI of the atlas, averaged by ROI.

We we used generalized psychophysiological interaction (gPPI) measures, representing the level of task-modulated (often labelled ‘effective’) connectivity between ROI or between ROI and voxels. gPPI is computed using a separate multiple regression model for each target (ROI). Each model includes three predictors: 1) task effects convolved with a canonical hemodynamic response function (psychological factor); 2) each seed ROI BOLD time series (physiological factor) and 3) the interaction term between the psychological and the physiological factors, the output of which is regression coefficients associated with this interaction term. Finally, group-level analyses were performed on these regression coefficients to assess for main effects within-group for contrasts of interest in seed-to-seed and seed-to-voxel analyses. Type I error was controlled by the use of seed-level FDR correction with *p*<.05 two tailed to correct for multiple comparison.

## Acknowledgements

The present study was supported by the Swiss National Science Foundation (SNSF 105314_146559/1) and the National Center for Affective Sciences (51NF40-104897). SF receives additional support from the SNSF (PP00P1_157409/1 and PP00P1_183711/1). LC helped program the tasks, collected the behavioral and neuroimaging data, analyzed the data, created and edited the figures and wrote the manuscript. MM collected TMS data and neuroimaging data and helped write the methods of the manuscript. DG helped design the study. CR helped design the study, especially the neuroimaging part to make it compatible with the TMS procedure. SF designed the study, programmed the tasks, collected part of the data and helped design the figures. All authors reviewed and edited the manuscript. The authors declare no competing interests whatsoever. The data and codes can be provided by Leonardo Ceravolo pending scientific review and a completed material transfer agreement. Requests for the data and codes should be submitted to.

## Notes

### Competing Interest Statement

The authors have declared no competing interest.

### Summary of Updates

Main text and some editing

