## Supplementary figures and tables for "Disrupting inferior frontal cortex activity alters affect decoding efficiency from clear but not from ambiguous affective speech"

**Table S1. Paired comparisons of stimulus morphing levels reaction times (initial stimulus evaluation).**

| Contrast | Degrees of freedom | Chi-square | *p*-value (Bonferroni) |
|---|---|---|---|
| $A_{100}$-$A_{90}$ | 1 | 184.18 | <.0001 |
| $A_{90}$-$A_{80}$ | 1 | 173.25 | <.0001 |
| $A_{80}$-$A_{70}$ | 1 | 191.18 | <.0001 |
| $A_{70}$-$A_{60}$ | 1 | 191.15 | <.0001 |
| $A_{60}$-$AF_{50}$ | 1 | 203.35 | <.0001 |
| $AF_{50}$-$F_{60}$ | 1 | 184.21 | <.0001 |
| $F_{60}$-$F_{70}$ | 1 | 170.10 | <.0001 |
| $F_{70}$-$F_{80}$ | 1 | 166.33 | <.0001 |
| $F_{80}$-$F_{90}$ | 1 | 169.89 | <.0001 |
| $F_{90}$-$F_{100}$ | 1 | 148.43 | <.0001 |

A: percentage of stimulus anger; F: percentage of stimulus fear; Test: Type II Wald Chi-square test.

**Table S2. Paired comparisons of stimulus morphing levels for initial stimulus evaluation.**

| Contrast | Degrees of freedom | Chi-square | *p*-value (Bonferroni) |
|---|---|---|---|
| $A_{100}$-$A_{90}$ | 1 | 227.55 | <.0001 |
| $A_{90}$-$A_{80}$ | 1 | 246.10 | <.0001 |
| $A_{80}$-$A_{70}$ | 1 | 266.27 | <.0001 |
| $A_{70}$-$A_{60}$ | 1 | 285.97 | <.0001 |
| $A_{60}$-$AF_{50}$ | 1 | 364.10 | <.0001 |
| $AF_{50}$-$F_{60}$ | 1 | 515.60 | <.0001 |
| $F_{60}$-$F_{70}$ | 1 | 553.39 | <.0001 |
| $F_{70}$-$F_{80}$ | 1 | 585.05 | <.0001 |
| $F_{80}$-$F_{90}$ | 1 | 597.67 | <.0001 |
| $F_{90}$-$F_{100}$ | 1 | 606.35 | <.0001 |

A: percentage of stimulus anger; F: percentage of stimulus fear; Test: Type II Wald Chi-square test.

**Table S3. Paired comparisons of reaction times as a function of voice morphing levels for the vocal affect categorization task.**

| Contrast | Degrees of freedom | Chi-square | *p*-value (Bonferroni) |
|---|---|---|---|
| $A_{90}$-$A_{70}$ | 1 | 3.84 | .75 |
| $A_{90}$-$AF_{50}$ | 1 | 5.85 | .23 |
| $A_{90}$-$F_{70}$ | 1 | 46.27 | <.0001 |
| $A_{90}$-$F_{90}$ | 1 | 67.72 | <.0001 |
| $F_{70}$-$AF_{50}$ | 1 | 0.21 | 1 |
| $F_{70}$-$F_{70}$ | 1 | 23.44 | <.0001 |
| $F_{70}$-$F_{90}$ | 1 | 39.28 | <.0001 |
| $AF_{50}$-$F_{70}$ | 1 | 19.22 | <.0001 |
| $AF_{50}$-$F_{90}$ | 1 | 33.77 | <.0001 |
| $F_{70}$-$F_{90}$ | 1 | 2.03 | 1 |

A: percentage of stimulus anger; F: percentage of stimulus fear; Test: Type II Wald Chi-square test.

**Table S4. Wholebrain results for the sample-specific voice areas functional localizer [vocal > non-vocal] contrast for the both groups together.**

| Region label | Hemisphere | MNI *X* | MNI *Y* | MNI *Z* | T-value | Cluster size (voxel count) |
|---|---|---|---|---|---|---|
| STS | R | 56 | -28 | 0 | 10.82 | 2874 |
| STG | L | -56 | -18 | -6 | 10.57 | 2501 |
| MFG | R | 42 | 6 | 34 | 5.05 | 1338 |
| IFGorb | L | -42 | 30 | -2 | 4.81 | 492 |

Statistical threshold: $p<.05$ FWE cluster correction (voxelwise $p<.005$ uncorrected, $k>59$).

*STS* superior temporal sulcus; *STG* superior temporal gyrus; *MFG* middle frontal gyrus; *IFGorb* inferior frontal gyrus *pars orbitalis*; L: left hemisphere; R: right hemisphere.
